# Sequence variation among SARS-CoV-2 isolates in Taiwan

**DOI:** 10.1101/2020.03.29.014290

**Authors:** Yu-Nong Gong, Kuo-Chien Tsao, Mei-Jen Hsiao, Chung-Guei Huang, Peng-Nien Huang, Po-Wei Huang, Kuo-Ming Lee, Yi-Chun Liu, Shu-Li Yang, Rei-Lin Kuo, Ming-Tsan Liu, Ji-Rong Yang, Cheng-Hsun Chiu, Cheng-Ta Yang, Shin-Ru Shih, Guang-Wu Chen

## Abstract

Taiwan experienced two waves of imported cases of coronavirus disease 2019 (COVID-19), first from China in January to late February, followed by those from other countries starting in early March. Additionally, several cases could not be traced to any imported cases and were suspected as sporadic local transmission. Twelve full viral genomes were determined in this study by Illumina sequencing either from virus isolates or directly from specimens, among which 5 originated from clustered infections. Phylogenetic tree analysis revealed that these sequences were in different clades, indicating that no major strain has been circulating in Taiwan. A deletion in open reading frame 8 was found in one isolate. Only a 4-nucleotide difference was observed among the 5 genomes from clustered infections.

## Introduction

A novel coronavirus emerged from Wuhan, Hubei province in China in December 2019 (1). This virus has been designated as severe acute respiratory syndrome coronavirus 2 (SARS-CoV-2), and the disease is named as coronavirus disease 2019 (COVID-19). The World Health Organization declared this disease a Public Health Emergency of International Concern on January 30, 2020. As of March 26, 2020, the outbreak of COVID-19 has resulted in 462,684 confirmed cases and 20,834 deaths worldwide (2), and 252 confirmed cases and two deaths were reported in Taiwan (3).

There have been two waves of COVID-19 cases in Taiwan. The first occurred from late January to the end of February, with most cases imported from China, either by Chinese tourists or Taiwanese businessmen returning for Chinese New Year. This wave was smaller than the second wave. The second wave started in early March, during which the disease occurred largely in Taiwanese tourists, business travelers, or students returning from other countries. Although most of these cases were traced to their foreign origins, some small and clustered infections were suspected to have been acquired by local transmission.

In this study, we performed virus culture and full-genome sequencing of isolates or clinical specimens of SARS-CoV-2. We compared the genomes obtained from Taiwanese samples to those of other strains in a database to understand their evolutionary trajectory. An open reading frame 8 (ORF8) deletion was found in one strain. Moreover, we assessed the number of nucleotide substitutions that may have accumulated in clustered infections during a short period of time.

## Methods

### Specimen Collection

Infection of patients by COVID-19 was confirmed by real-time reverse-transcriptase polymerase chain-reaction (RT-PCR) according to the guidelines of the Taiwan Centers for Disease Control (CDC; https://www.cdc.gov.tw/En), and all nasopharyngeal (NP), throat (TH) swab, and sputum (SP) samples were maintained in universal transport medium for further analysis.

### Cell Culture and Virus Isolation

Vero-E6 (ATCC, Manassas, VA, USA) and MK-2 (ATCC) cells were maintained in Modified Eagle Medium (MEM, Thermo Fisher Scientific, Waltham, MA, USA) supplemented with 10% fetal bovine serum and 1x penicillin-streptomycin at 37°C in the presence of 5% CO_2_. To isolate the virus, all procedures following the laboratory biosafety guidelines of the Taiwan CDC were conducted in a biosafety Level-3 facility. Cells grown to 80–90% confluency in a T-25 flask were inoculated with 500 μL of virus solution, which was prepared by diluting 100 μL of specimen samples with 1.5 mL of sample pretreatment medium consisting of MEM and 2× penicillin-streptomycin solution, followed by incubation at 37°C for 1 h. The absorption was performed at 37°C for 1 h, then cells were refreshed with 5 mL virus culture medium composed of MEM, 2% fetal bovine serum, and 1× penicillin-streptomycin solution and maintained at 37°C. Infected cells were observed daily to determine their cytopathic effect. Additionally, RT-PCR analysis using the RNA extracted from part of the culture supernatant every two days after inoculation was performed to monitor viral growth. We continuously observed the infected cells until cytopathic effects occurred in more than 75% of the cells, after which the culture supernatant was harvested.

### Whole-Genome Sequencing

RNA was extracted either from the culture supernatant or directly from the specimens using a QIAmp viral RNA mini Kit (Qiagen, Hilden, Germany) according to the manufacturer’s instructions, except that the carrier RNA was replaced with linear acrylamide (Thermo Fisher Scientific) as the co-precipitant. The amount of viral RNA was evaluated by quantitative RT-PCR to examine the Ct value of the viral E gene. For RNAs showing a high Ct value, we used the Ovation RNA-Seq System V2 (Nugen Technologies, San Carlos, CA, USA) to synthesize cDNA which was further processed into a library using the Celero DNA-Seq System (Nugen Technologies). Other samples with lower Ct values were used for library preparation by using the Trio RNA-Seq kit (Nugen Technologies). Sequencing was performed on an Illumina MiSeq System (San Diego, CA, USA) with paired-end reads. More than 0.75 and 2.5 Gb of raw data were generated for samples from viral isolates and clinical specimens, respectively.

### Next-generation Sequencing Data Analysis Pipeline

We first trimmed the raw data by removing low-quality and short reads using Trimmomatic (version 0.39) (4). Next, quality reads were mapped to the human reference genome to remove host sequences using HISAT2 (version 2.1.0) (5). SPAdes (version 3.14.0) (6) was used to perform *de novo* assembly for constructing contig sequences. Fourth, the BLASTN tool was used to search the assembled contigs against the nucleotide sequence (NT) database of the National Center for Biotechnology Information (NCBI). Viral candidates were identified using the reported top BLASTN hits for each of the queried contig sequences. Finally, we used an iterative mapping approach (7) to increase the read depth and coverage of quality contigs to obtain the whole genome.

### Phylogenetic and Sequence Analysis

Twelve whole genomes were assembled by using our pipeline, including three genomes from specimens and nine genomes from isolates, which were deposited in the Global Initiative on Sharing All Influenza Data (GISAID, https://www.gisaid.org/) with accession numbers EPI_ISL_411915, EPI_ISL_417518, EPI_ISL_415741–3, and EPI_ISL_417519–25, according to CGMH-CGU No. 1–12. We further downloaded all complete and high-coverage genomes from GISAID as of March 14, 2020, and obtained 335 sequences after removing those with sequences gaps or ambiguous nucleotides. One reference strain (accession number MN908947.3) was downloaded from GenBank (NCBI). In total 348 sequences were aligned using MAFFT (version 7.427) (8) for further analyses. The phylogenetic tree was inferred using RAxML (version 8.2.12) (9) under the GTRGAMMA model with a bootstrap value of 1000 to investigate the genomic relationships.

## Results

### Phylogenetic Tree of Taiwanese and Global Strains

Twelve complete genomes from three specimens (CGMH-CGU No. 1, 7, and 8) and nine isolates (No. 2–6 and 9–12) were uploaded to GISAID. Table 1 shows their next-generation sequencing (NGS) coverage and depth. All average depths were greater than 10,000, except for CGMH-CGU-04 and −08 which showed values of 446.0 and 53.0, respectively. Table 1 also includes two earlier strains, hCoV-19/Taiwan/2/2020 and hCoV-19/Taiwan/3/2020, previously submitted by Taiwan CDC.

**Table 1.** Specimen collection, culture, and sequencing.

| **CGMH-CGU**<br>**ID/Strain name \*\*** | **Collection**<br>**date** | **Viral culture**<br>**(day)** | **Source\* (Ct value of E**<br>**gene)** | **Coverage and**<br>**avg. depth of**<br>**SARS-CoV-2** |
| --- | --- | --- | --- | --- |
| 1 | 2020-01-25 | - | SP (17.01) | 99.9%; 1157.4 |
| 2 | 2020-02-04 | 14 | MK2 (10.0) | 100.0%; 4735.8 |
| 2 | 2020-02-04 | - | NP (29.07) | 80.0%; 3.3 |
| 3 | 2020-02-26 | 10 | MK2 (14.25) | 100.0%; 18,299.0 |
| 4 | 2020-02-27 | 4 | Vero E6 (26.15) | 99.2%; 446.0 |
| 5 | 2020-02-27 | 4 | MK2 (12.78) | 100.0%; 26,521.5 |
| 6 | 2020-03-05 | 5 | MK2 (12.82) | 100.0%; 13,029.9 |
| 7 | 2020-03-09 | - | SP (22.98) | 99.9%; 53.0 |
| 8 | 2020-03-10 | - | NP (23.18) | 100.0%; 10,412.2 |
| 9 | 2020-03-13 | 3 | MK2 (10.89) | 100.0%; 30,044.7 |
| 10 | 2020-03-13 | 3 | MK2 (10.45) | 100.0%; 29,614.0 |
| 11 | 2020-03-14 | 3 | Vero E6 (11.08) | 100.0%; 24,326.9 |
| 12 | 2020-03-14 | 3 | MK2 (10.11) | 100.0%; 34,422.0 |
| TW/2 | 2020-01-23 | - | - | - |
| TW/3 | 2020-01-24 | - | - | - |

\* Sources from sputum (SP), nasopharyngeal swab (NP), and throat swab (TH) specimens, or supernatant on MK2 and Vero E6 cells
\*\*\* Twelve GISAID accession numbers of CGMH-CGU No. 1–12 are EPI\_ISL\_411915, EPI\_ISL\_417518, EPI\_ISL\_415741–3, and EPI\_ISL\_417519 –25. The other two Taiwanese
strains (TW/2 and TW/3) were previously submitted to GISAID by Taiwan CDC, with accession numbers (EPI\_ISL\_406031 and EPI\_ISL\_411926, respectively).

The phylogenetic tree revealed that the SARS-CoV-2 viral genomes from Taiwan (highlighted) were in different clades (Figure 1). Viral genomes of No. 3–7 were from clustered infections, together with No. 8 (a case originating from the United Kingdom), and some from Australia (AUS) and New Zealand (NZ) in the yellow clade. Three patients with AUS/NZ infections had a travel history to Iran. This figure also shows eight additional Taiwanese isolates (highlighted), which appeared in distinct lineages, indicating that no single dominant strain has been circulating in Taiwan.

**Figure 1.**
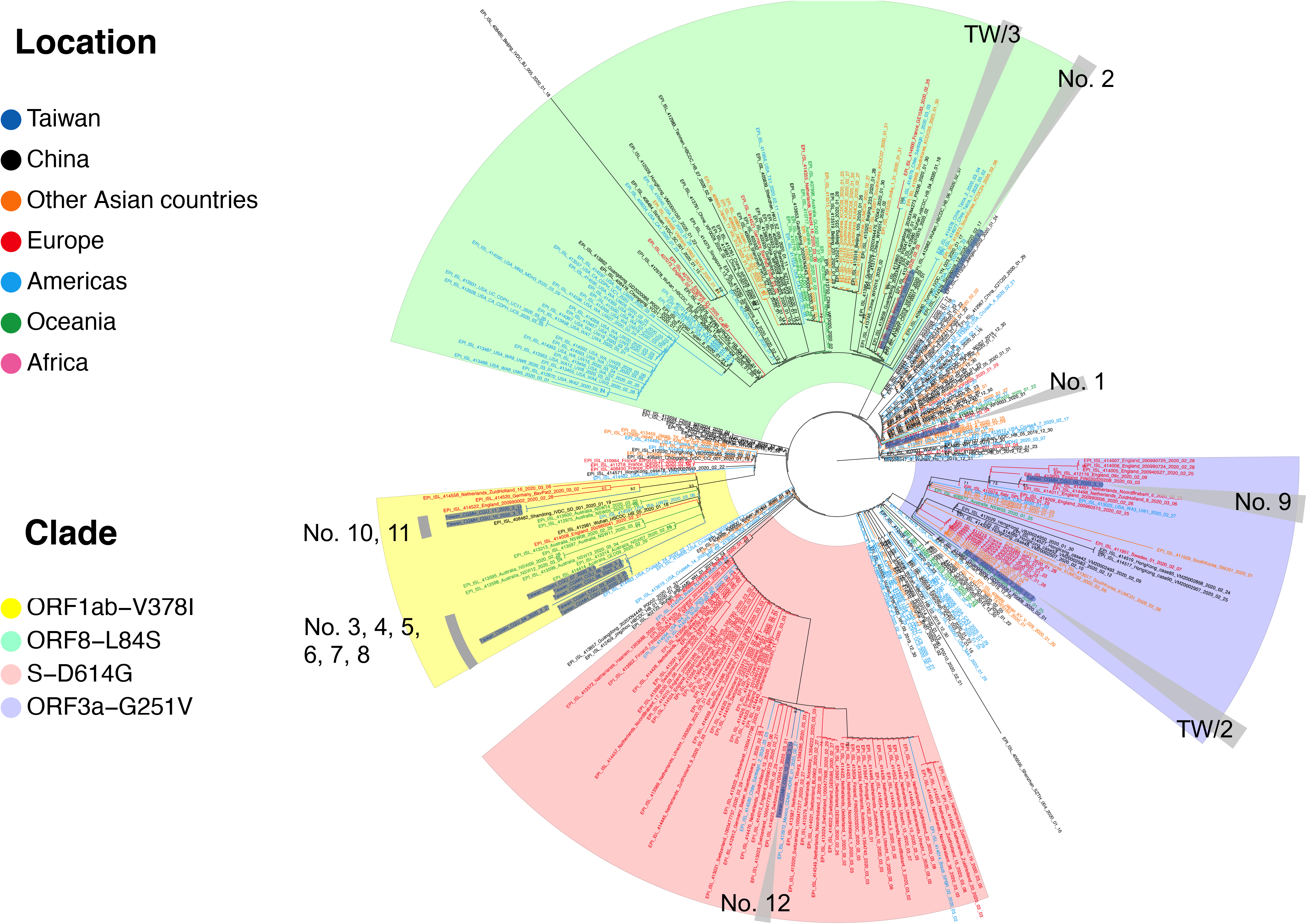
Phylogenetic tree of Taiwanese and global strains. Phylogeny was inferred using a maximum likelihood approach. Taiwanese strains are highlighted. Strains isolated from different locations and clades with specific variations are marked in different colors. Significant bootstrap support values greater than 70% are shown.

The two earliest sequences in this yellow clade were dated to mid-January from Wuhan and Shangdong, which may have been the origin of the yellow clade. CGMH-CGU-03 had no travel history and the specimen was collected nearly 6 weeks after the two Chinese isolates were collected. All other viruses in this clade were also dated after February 26. Separated by this long duration from the two Chinese strains in mid-January, it is unlikely the later strains were directly linked to the Wuhan strains. Although some AUS/NZ cases in this clade had a travel history to Iran, the transmission route of these five Taiwanese cases remains unclear.

### ORF8 Deletion Revealed by NGS Data Analysis

Figure 2A shows the NGS coverage and depth of CGMH-CGU-01. This strain was identical to the WuHan-1 strain (accession number MN908947.3). The most divergent strain among the 14 Taiwanese sequences was CGMH-CGU-04 which showed nine nucleotide changes (resulting in five amino acid changes) in the coding region compared to CGMH-CGU-01. Notably, we detected a deletion in a 382-nucleotide (nt) sequence at genomic positions 27,848– 28,229 in CGMH-CGU-02. Figure 2B shows the coverage and depth of this strain. According to the reference strain (WuHan-1), the genomic position of ORF8 was 27,894–28,259 (Figure 2C). This 382-nt deletion begins upstream of ORF8 to nearly the end of ORF8. We further performed NGS using a specimen isolated from the same patient. Reads yielding this 382-nt deletion were confirmed in original specimen, although only the partial genome was assembled (Table 1).

**Figure 2.**
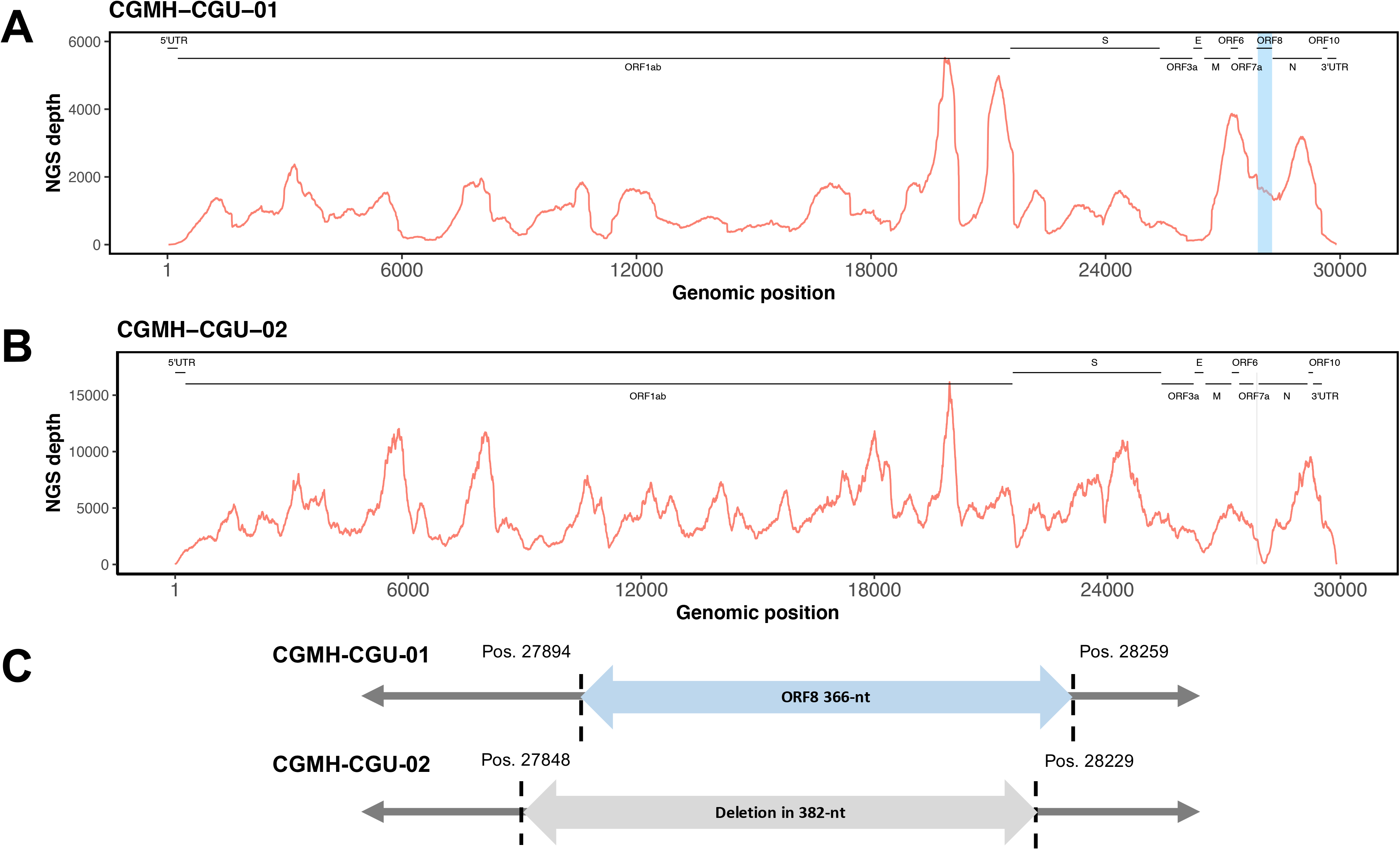
ORF8 deletion in SARS-CoV-2 genome. A and B) NGS depths of CGMH-CGU-01 and CGMH-CGU-02 and C) genomic regions of ORF8 and ORF8 deletion according to the reference strain are shown.

### Within Four Nucleotide Changes among Virus Isolates from Clustered Patients

COVID-19 has been reported to be transmitted through close contact among confirmed cases. Regardless of whether individuals are symptomatic, their family members and co-workers are at risk of becoming infected. Viral genomes No. 4–7 were from patients who had contact with an index patient (CGMH-CGU-03). To identify the number of nucleotides changed in the viral genome during clustered infections, we determined the viral full genomes either from viral isolates (No. 3–6) or specimens (No. 7) of these 5 cases. Although the genomes of samples No. 3, 5, and 6 were identical, they differed from that of No. 4 at 3 ORF1ab nucleotide positions A4788G, C10809T, and G21055A; the third position showed a synonymous change with a G7019S amino acid substitution (Figure 3). Number 7 showed only one nucleotide difference from No. 3, 5, and 6. These results suggest that only 4 nucleotide changes occurred in the viral genome among cases in clustered infections.

**Figure 3.**
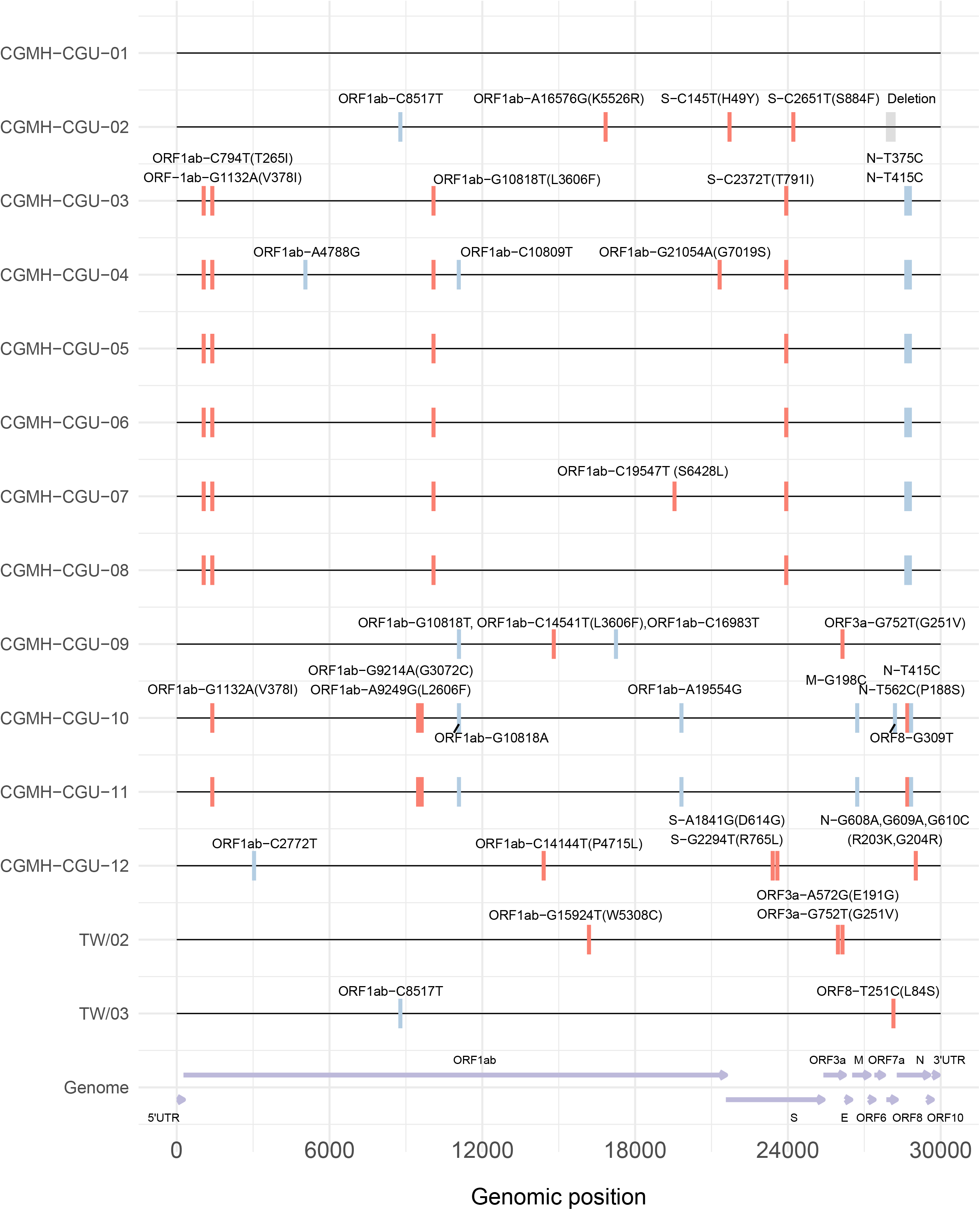
Nucleotide and amino acid variations in SARS-CoV-2 genomes. Compared to CGMH-CGU-01 (identical to the reference strain), nucleotide and amino acid variations in the SARS-CoV-2 genomes from Taiwanese strains are shown. Synonymous and nonsynonymous mutations were marked by blue and red color, respectively. Amino acid changes were annotated in parentheses. ORF8 deletion was marked in gray.

## Discussion

Twelve full viral genomes were resolved in this study either from virus isolates or directly from specimens. Phylogenetic tree analysis showed that these sequences were in different clades, indicating that no major strain is currently circulating in Taiwan. A deletion in ORF8 was found in one isolate, which has also been detected in patients in Singapore (10). Four or fewer nucleotide differences were observed in the 5 genomes from clustered infections.

We detected a 382-nt deletion covering nearly the entire open reading frame 8 of the CGMH-CGU-2 isolate obtained from a patient who returned from Wuhan in January. A similar observation was reported for eight hospitalized patients in Singapore. During the SARS-CoV outbreak in 2003, deletions in ORF8 were observed, which were associated with a reduced ability for virus replication in human cells (11).

RNA viruses show variations in their genomes due to nucleotide substitutions generated by the low fidelity of RNA-dependent RNA polymerase during replication. The genome variation of these viruses is thought to facilitate successful adaption to the environment of various hosts. However, previous studies showed that the mutation rates of RNA viruses vary in different viruses and depend on the viral transmission modes (12). Sequence analysis of SARS-CoV-2 isolated from 5 patients from February 26 to March 9, 2020 in CGMH Taiwan revealed only 4 mutations in their 29,903-nt genomic RNA. This suggests that the nucleotide substitution rate is controlled during viral RNA replication. The nsp14 exoribonuclease encoded by several coronaviruses plays a role in proofreading during genome replication (13, 14); further studies are required to investigate the function of SARS-CoV-2 nsp14 in replication fidelity.

Timely sharing full genomes of SASR-CoV-2 from different locations is important for monitoring genetic changes in the virus which may be associated with viral spreading and clinical manifestations. We determined the sequences of SARS-CoV-2 in Taiwan in different clades. Moreover, four or fewer nucleotide changes in viral genomes from five cases in clustered infections indicated that sequencing is a useful tool for tracing the source of infection for this type of RNA virus.

## Acknowledgments

This work was financially supported by the Research Center for Emerging Viral Infections from The Featured Areas Research Center Program within the framework of the Higher Education Sprout Project by the Ministry of Education (MOE) in Taiwan, the Ministry of Science and Technology (MOST), Taiwan (MOST 108-3017-F-182-001, MOST 107-2221-E-182-064-MY2, and MOST 106-2320-B-182A-013-MY3), and Linkou Chang Gung Memorial Hospital, Taiwan (No. CLRPG3B0048 and CMRPD1H0231–3).

